# Genome-wide association studies of impulsive personality traits (BIS-11 and UPPSP) and drug Experimentation in up to 22,861 adult research participants

**DOI:** 10.1101/414854

**Authors:** Sandra Sanchez-Roige, Pierre Fontanillas, Sarah L. Elson, the 23andMe Research Team, Michelle Agee, Babak Alipanahi, Adam Auton, Robert K. Bell, Katarzyna Bryc, Sarah L. Elson, Pierre Fontanillas, Nicholas A. Furlotte, David A. Hinds, Karen E. Huber, Aaron Kleinman, Nadia K. Litterman, Jennifer C. McCreight, Matthew H. McIntyre, Joanna L. Mountain, Elizabeth S. Noblin, Carrie A.M. Northover, Steven J. Pitts, J. Fah Sathirapongsasuti, Olga V. Sazonova, Janie F. Shelton, Suyash Shringarpure, Chao Tian, Joyce Y. Tung, Vladimir Vacic, H. Catherine, Joshua C. Gray, Harriet de Wit, James MacKillop, Abraham A. Palmer

**Author notes:** Correspondence: Professor AA Palmer, Department of Psychiatry, University of California San Diego, 9500 Gilman Drive, Mail Code 0667, BRF2 3A24, La Jolla, CA 92093, USA. Collaborator List for the 23andMe Research Team.

## Abstract

**Background:** Impulsive personality traits are complex heritable traits that are governed by frontal-subcortical circuits and are associated with numerous neuropsychiatric disorders, particularly drug abuse.

**Methods:** In collaboration with the genetics company 23andMe, Inc., we performed several genome-wide association studies (**GWAS**) on measures of impulsive personality traits (the short version of the UPPSP Impulsive Behavior Scale, and the Barratt Impulsiveness Scale [BIS-11]) and drug experimentation (the number of drug classes an individual has tried in their lifetime) in up to 22,861 male and female adult research participants of European ancestry.

**Results:** Impulsive personality traits and drug experimentation showed SNP-heritabilities that ranged from 5 to 11%. Genetic variants in the *CADM2* locus were significantly associated with the UPPSP Sensation Seeking subscale (*P* = 8.3 × 10^-9^, rs139528938) and showed a suggestive association with drug experimentation (*P* = 3.0 × 10^-7^, rs2163971; r^2^ = 0.68 with rs139528938); *CADM2* has been previously associated with measures of risky behaviors and self-reported risk tolerance, cannabis initiation, alcohol consumption, as well as information speed processing, body mass index (BMI) variation and obesity. Furthermore, genetic variants in the *CACNA1I* locus were significantly associated with the UPPSP Negative Urgency subscale (*P* = 3.8 × 10^-8^, rs199694726). Multiple subscales from both UPPSP and BIS showed strong genetic correlations (>0.5) with drug experimentation and other substance use traits measured in independent cohorts, including smoking initiation, and lifetime cannabis use. Several UPPSP and BIS subscales were genetically correlated with attention-deficit/hyperactivity disorder (r_g_ = 0.30-0.51, p < 8.69 x 10^-3^), supporting their validity as endophenotypes.

**Conclusions:** Our findings demonstrate a role for common genetic contributions to individual differences in impulsivity. Furthermore, our study is the first to provide a genetic dissection of the relationship between different types of impulsive personality traits and various psychiatric disorders.

---

Impulsive personality traits (**IPTs**) are complex traits (1–6) that are associated with the risk for several major neuropsychiatric disorders, including substance use disorders. In addition, IPTs influence temporally distinct stages of drug abuse, from experimentation to dependence (7, 8).

Self-report questionnaires are commonly used to quantify IPTs in normal adults and patient populations. The Impulsive Behavior Scale, **UPPSP** (9, 10), and the Barratt Impulsiveness scale, **BIS-11** (11), are arguably the two most common questionnaires. Despite evidence from twin and family studies showing that IPTs are moderately heritable [40-60% (12–16)], it has proven difficult to identify specific genes that influence impulsivity (17, 18). Candidate gene studies have implicated several genetic loci with UPPSP and BIS scores (see (19) for a list of previously associated *a priori* loci), but those results have generally failed to replicate. The largest prior study of the genetic basis of the trait impulsivity in the UPPSP and BIS used 983 healthy young individuals of European ancestry (19) and was thus underpowered.

In collaboration with the personal genetics company 23andMe, Inc., we performed the largest GWASs of IPTs to date using scores from the UPPSP and BIS-11 questionnaires in up to 22,861 research participants. IPTs, although conceptually related, are believed to be independent constructs (20, 21), with distinct underlying neurobiological and neurochemical substrates (3, 22). Therefore, we hypothesized that different facets of impulsivity among the UPPSP and BIS-11 would show both a modest overlap and distinct genetic architecture. Furthermore, different IPT dimensions may have a different role in various forms of psychopathology. For example, different facets of impulsivity may influence different stages of drug abuse, including experimentation, acquisition of regular drug use, progression to addiction, failure to quit, and risk of relapse (23–25). To directly examine differential genetic influences on impulsivity that may contribute to early stages of drug use initiation, we also performed a GWAS on level of drug experimentation, which quantifies the number of drug classes that an individual has tried in their lifetime. These GWASs were then extended with gene- and transcriptome-based analyses. Finally, we examined the genetic correlation among the IPTs and drug experimentation, as well as genetic correlations with related phenotypes in archival datasets.

## MATERIALS AND METHODS

### Sample

All participants included in the analyses were drawn from the customer base of 23andMe, Inc., a consumer genetics company. Participants provided informed consent and answered surveys online under a protocol approved by Ethical and Independent Review Services, an independent AAHRPP-accredited institutional review board (http://www.eandireview.com). We restricted our analysis to unrelated participants of European ancestry (>97% as determined through an analysis of local ancestry (26); see **Supplementary** for additional details) for whom UPPSP, BIS and Drug Experimentation data were available. The final number of research participants included in the analyses range from 21,495 to 22,861. Recruitment occurred over an approximately four-month period in 2015. This sample has been extensively described elsewhere (27, 28). Sociodemographic details are described in the **Supplementary Table 1**.

### Self-reported impulsivity traits

To measure IPTs we used four subscales from the 20-item UPPSP Impulsive Behavior Scale [brief version, **UPPSP**; (9, 10)]: (lack of) Premeditation, (lack of) Perseverance, Positive Urgency and Negative Urgency, Sensation Seeking. Each subscale includes 4-items and yields scores from 4 to 16. We also administered the Barratt Impulsiveness Scale [**BIS-11**; (11)], a 30-item measure that gives a total impulsivity score and 3 subscores of Attentional, Motor, and Nonplanning impulsiveness. We used UPPSP and BIS because they represent the most commonly used multifaceted measures of IPTs. Since the scores were not normally distributed across these measures (by visual inspection, **Supplementary Figures 1-2**, **Supplementary Tables 1-2**), we used a quantile normalization prior to GWAS analyses.

### Drug Experimentation

The measure of drug experimentation quantifies the number of 11 different classes of drugs an individual has used (tobacco [cigarettes, cigars, chewing tobacco], alcohol, marijuana, cocaine, methamphetamine, LSD/magic mushrooms, ecstasy, prescription stimulants [taken not as prescribed; e.g., Ritalin, Adderall, Strattera], prescription painkillers [taken not as prescribed; e.g., Vicodin, Oxycontin], heroin, opium); this measure yields scores from 0 to 11, and was adapted from the PhenX toolkit (www.phenxtoolkit.org/). We used a quantile normalization, since scores were not normally distributed (by visual inspection, **Supplementary Figure 3**, **Supplementary Table 3**).

### Genotyping, quality control and imputation

We have previously reported a full description of these methods (27). DNA extraction and genotyping were performed on saliva samples by CLIA-certified and CAP-accredited clinical laboratories of Laboratory Corporation of America. Samples were genotyped on 23andMe custom genotyping array platforms (Illumina HumanHap550+ Bead chip V1 V2, OmniExpress+ Bead chip V3, Custom array V4). Quality control of genetic variants and imputation were performed by 23andMe (see **Supplementary Table 4**).

### SNP-heritability using LD Score Regression

We used Linkage Disequilibrium Score Regression Coefficient [LDSC (29)] to measure SNP-heritability of Sensation Seeking and drug experimentation. To standardize the input file (GWAS summary statistics), we followed quality controls as implemented by the LDSC python software package. We used pre-calculated LD scores [“eur_w_ld_chr/” files (30); MHC region excluded] for each SNP using individuals of European ancestry from the 1000 Genomes project, suitable for LD score analysis in European populations. We restricted the analysis to well-imputed SNPs: the SNPs were filtered to HapMap3 SNPs (31), and were required to have a minor allele frequency (MAF) above 1%. InDels, structural variants, strand-ambiguous SNPs, and SNPs with extremely large effect sizes (᙭^2^ > 80) were removed. One of the advantages of using LDSC is that it allowed us to distinguish between genomic inflation attributed to polygenic architecture, from confounding biases such as population stratification (29). As expected under polygenicity, we observed inflation of the test statistic (Mean ᙭^2^ < 1.05), and adjusted for a genomic control inflation factor λ (the ratio of the observed median ᙭^2^ to that expected by chance). LD score intercepts of less than 1.01 (SE = 0.01) suggested that deviation from the null was due to a polygenic structure rather than inflation due to population structure biases (see **Supplementary Table 5**).

### Genome-wide association analyses

GWAS analyses were performed using the 23andMe internal pipeline, which we have previously described (27). We performed association tests by linear regression (additive model). We included age (inverse-normal transformed), sex, the first five principal components of genotype, and indicator variables for genotype platforms as covariates (**Supplementary Table 6**). Additional details can be found in the **Supplementary Material**.

### Gene-based and transcriptome-based analyses

We performed MAGMA (32) competitive gene-set and pathway analyses using the summary statistics from the GWAS of UPPS and BIS subscales, and drug experimentation using FUMA v1.2.8 (32). SNPs were mapped to 18,133 protein-coding genes from Ensembl build 85. Gene-sets were obtained from Msigdb v5.2 (“Curated gene sets”, “GO terms”).

We used S-PrediXcan (33) to predict gene expression levels in 10 brain tissues, and to test whether the predicted gene expression correlates with GWAS of UPPSP, BIS and drug experimentation. We used pre-computed tissue weights from the Genotype-Tissue Expression (**GTEx** v6) project database (www.gtexportal.org/) as the reference transcriptome dataset. Further details are provided elsewhere (27).

### Phenotypic and genetic correlation analyses

We used bivariate correlations (*r*) to examine the direct phenotypic correlations between UPPSP, BIS, Drug Experimentation and several variables of interest (age, gender, race, education, annual household), and to identify significant covariates for inclusion in GWAS analysis, and inter-correlations between the UPPSP and BIS subscales (**Supplementary Tables 7-9**).

Using LDSC, we calculated genetic correlations (*r*_g_) between the 5 UPPSP subscales, 4 BIS traits and Drug Experimentation and 45 other complex traits or diseases that have been previously associated with IPTs, for which we had access to summary statistics. References for the datasets used are identified in **Supplementary Tables 10-19**. Files were standardized using the steps described in the section above (“SNP-heritability using LD Score Regression”). We did not constrain the intercepts in our analysis because the degree of sample overlap was unknown. We used False Discovery Rate (**FDR**) to correct for multiple testing (34).

## RESULTS

### Demographics

Demographic data are shown in **Table 1**. Mean age was 53.8 years (SD = 16.1), and 55.3% were women. The annual household income ranged from less than $14,999 (13.5%) to greater than $75,000 (21.5%), and the mean years of education completed was 16.8 (SD = 2.6), which is the equivalent of completing a bachelor degree. About half of the participants (49.3%) were married/partnered. Participants exhibited low to moderate alcohol and drug use (**Table 1**).

### Self-reported impulsivity and drug experimentation scores

Self-reported impulsivity and drug experimentation scores are shown in the **Supplementary Tables 1-3** and

**Supplementary Figures 1-3**.

### Chip-heritability estimates

We estimated chip-heritability (with LDSC), and demonstrated that 4.5 to 11.2% of the variation of UPPSP, BIS and drug experimentation can be explained by single-nucleotide polymorphisms. Full results are shown in the **Supplementary Table 5**.

### Genome-Wide Association Analyses of UPPSP, BIS and Drug Experimentation

The Manhattan and quantile-quantile (**QQ**) plots for UPPSP Sensation Seeking are shown in **Figure 1** and **Supplementary Figures 4 and 14**. We detected one genome-wide significant hit on Chromosome 3 (P = 8.3 × 10^-9^, rs139528938; **Supplementary Table 22**), located in the gene *CADM2* (Cell Adhesion Molecule 2), which encodes a member of the synaptic cell adhesion molecule 1 (SynCAM) family and belongs to the immunoglobulin (Ig) superfamily. *CADM2* has been recently associated with other risk-related phenotypes including risk-taking personality (35, 36) and risky behavior (37), alcohol consumption (38, 39) and cannabis use (40, 41), as well as being associated with information speed processing (42), physical activity (43) and BMI variation (44, 45). We used FUMA to functionally annotate all 467 SNPs in the credible set (see **Supplementary Table 23**). All SNPs were intronic. Furthermore, 18 SNPs showed CADD scores >12.37, which is the suggested threshold to be considered deleterious (46). Four SNPs had RegulomeDB scores of 1a-1f, showing evidence of potential regulatory effects. 59.7% of the SNPs were in open chromatin regions (minimum chromatin state 1-7).

**Figure 1.**
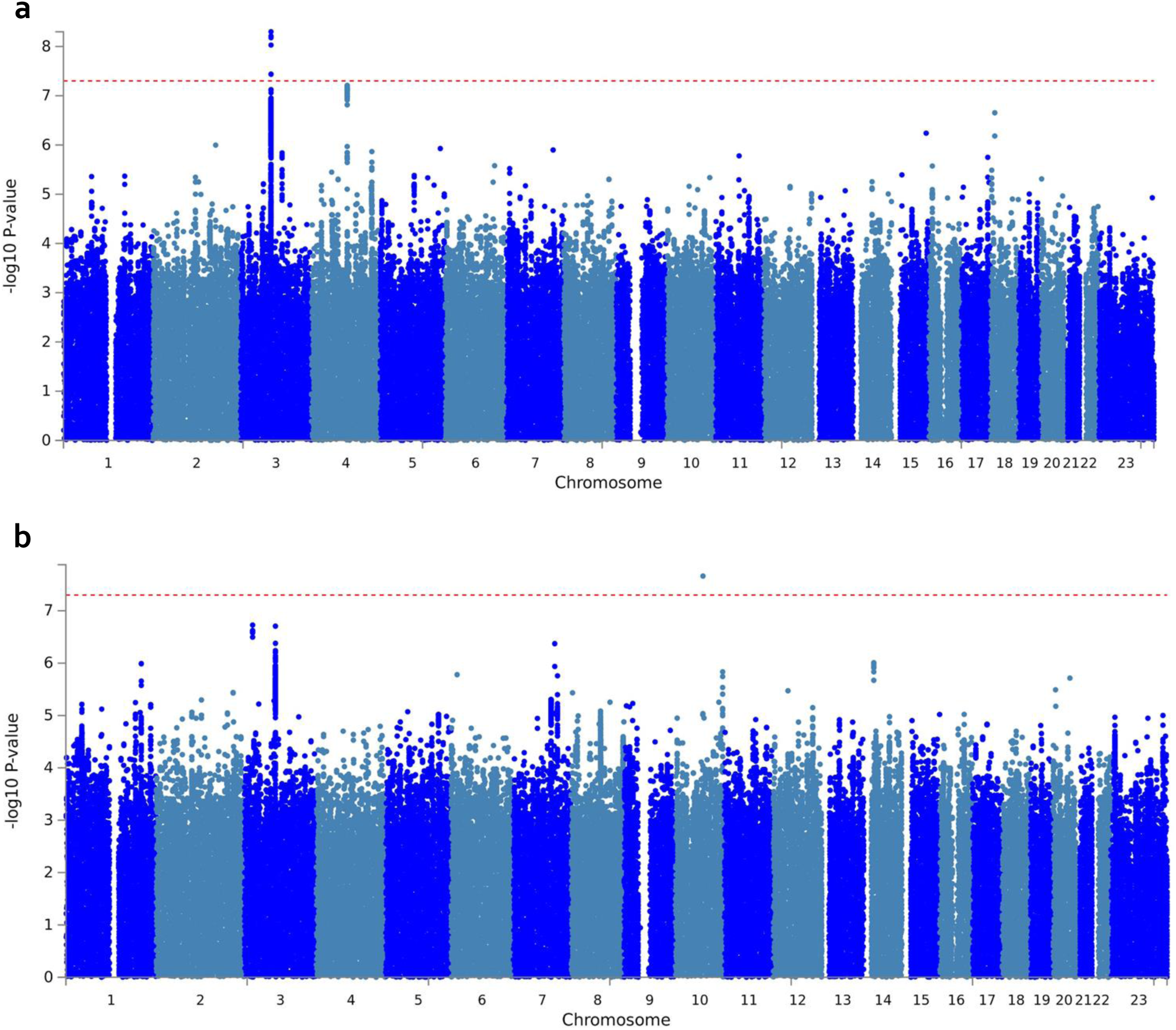
Manhattan plot of GWAS results indicating the strongest associations between the 22 autosomes, X chromosome, and Sensation Seeking (**a**) and drug experimentation (**b**). Line denotes genome-wide significance (p < 5 x 10^-8^). The results have been adjusted for a genomic control inflation factor λ=1.029 and λ=1.031 (sample size = 22,745 and 22,572, respectively).

In addition, we detected a suggestive association between drug experimentation and an intronic variant of *CADM2* (P = 3.0 × 10^-7^, rs2163971; **Figure 1**; **Supplementary Figures 13** and **23**). We also identified a novel association between genetic variants in the *CACNA1I* locus and the UPPSP Negative Urgency subscale (*P* = 3.8 × 10^-8^, rs199694726); the most associated SNP in this locus, rs4522708 (P = 8.22 x 10^-8^), is in LD (r^2^ = 0.64) with rs5995756, which has been previously associated with schizophrenia (**Figure 2**; **Supplementary Figures 7** and **17**)(47). We did not identify any associations that exceeded 5 x 10^-8^ for any of the other UPPSP and BIS subscales (**Supplementary Figures 5-12** for Manhattan and QQ plots; **Supplementary Table 22** and **Supplementary Figures 15-22** for the list of genetic variants and regional plots with strongest associations).

**Figure 2.**
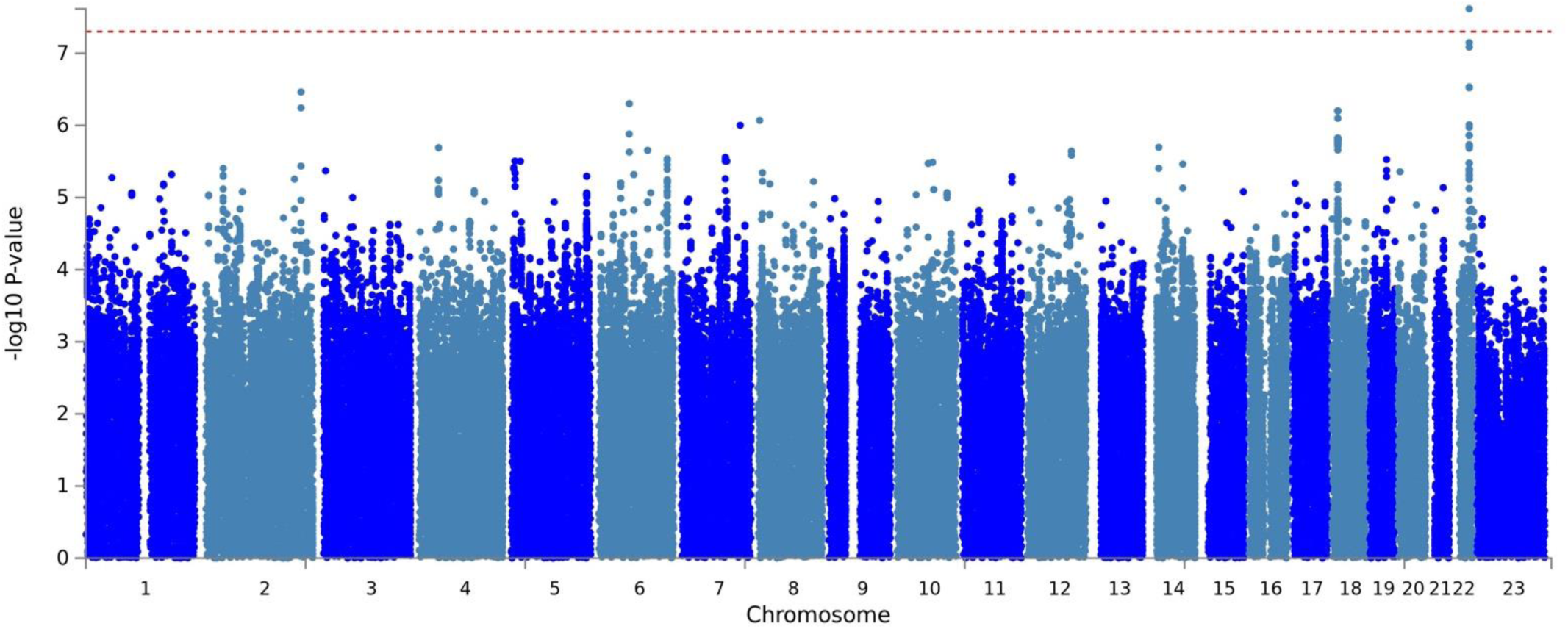
Manhattan plot of GWAS results indicating the strongest associations between the 22 autosomes, X chromosome, and UPPSP negative urgency. The results have been adjusted for a genomic control inflation factor λ=1.030 (sample size = 22,795).

### Gene and transcriptome-based analyses

Similar to the GWAS results, MAGMA also identified the gene *CADM2* as being significantly associated with Sensation Seeking (p = 1.7 x 10^-9^; Bonferroni threshold: p = 0.05/18,909 = 2.6 x 10^-6^; **Supplementary Figure 24**) and drug experimentation (p = 1.7 x 10^-7^; **Supplementary Figure 24b**). *CACNA1I* was associated with positive and Negative Urgency (**Supplementary Tables 27** and **28**). Finally, *MSRA* was associated with Negative Urgency [1.3 x 10^-6^; **Supplementary Table 28**; (49)]. MAGMA did not identify any canonical pathways that were significantly associated with any of the other IPTs studied (data not shown). Similar to the GWAS and MAGMA results, S-PrediXcan also identified a positive correlation (p = 3.8 x 10^-6^; FDR 20%) between Sensation Seeking and *CADM2* expression in the putamen (basal ganglia) (**Supplementary Table 34**). S-PrediXcan did not detect any other significant association for other UPPSP and BIS subscales, or drug experimentation (**Supplementary Tables 35-42**).

### Phenotypic and genetic correlations

A *phenotypic* correlation matrix of the impulsive personality subscales and drug experimentation is shown in the **Supplementary Tables 7-9**. All subscales exhibited adequate internal reliability (**Table 1**). BIS, UPPSP and drug experimentation scores were associated with demographic variables (age, sex, body mass index, household income, and years of education); male and younger research participants were more impulsive across all of the UPPSP and BIS subscales, and showed higher drug experimentation scores than female and older participants. Research participants with higher BMI and lower household income and years of education showed greater impulsivity scores as measured by UPPSP and BIS; such participants also showed higher scores for drug experimentation.

Consistent with recent observations (19, 20), phenotypic inter-correlations for UPPSP and BIS subscales were high and positive (except between UPPSP Sensation Seeking and UPPSP Perseverance; see **Supplementary Tables 7** and **8**). Across impulsivity questionnaires, phenotypic correlations between UPPSP and BIS subscales were also high and positive, except between UPPSP Sensation Seeking and BIS Nonplanning (*r* =-0.01. p = 5.3 × 10^-2^). Phenotypic correlations between Sensation Seeking and other impulsivity traits, although significantly different from zero, were modest (*r* <0.2). This is consistent with recent phenotypic modelling of latent interrelationships among indicators of impulsivity (20).

**Figure 3** shows a *genetic* correlation matrix of the impulsive personality subscales and several other phenotypes (full results shown in **Supplementary Tables 10-19**). Phenotypic and genetic correlations between impulsive subscales exhibited substantial variability (**Figure 4**, **Supplementary Tables 20-21**). All BIS subscales were highly genetically inter-correlated (r_g_ = 0.59-1, ps < 5.0 x 10^-9^). With regards to UPPSP subscales, UPPSP Positive Urgency showed a highly positive genetic correlation with Negative Urgency (r_g_ = 0.74, p = 1.60 x 10^-9^) and Premeditation (r_g_ = 0.62, p = 2.10 x 10^-3^), but we did not observe any significant genetic associations (FDR 5%) between the remaining three UPPSP subscales. Consistent with the observed phenotypic correlations, we did not find any significant genetic correlations between Sensation Seeking and other UPPSP or BIS subscales (r_g_ = -0.03-0.42, p < 7.85 x 10^-2^), suggesting that this trait may represent a relatively different construct, as previously suggested (20).

**Figure 3.**
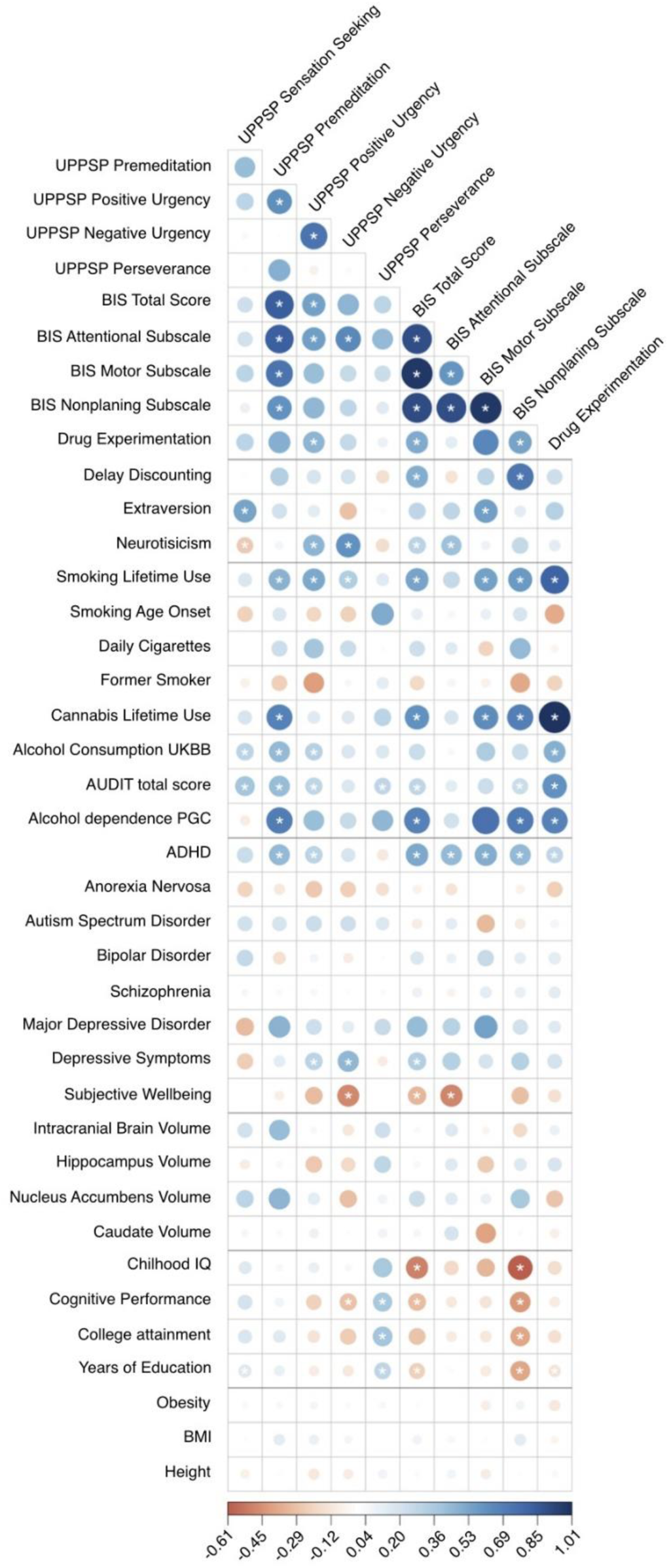
Genetic correlations between UPPSP, BIS, and drug experimentation and several traits: other impulsive and personality traits, substance use phenotypes, neuropsychiatric, brain volume, cognition, anthropomorphic.

**Figure 4.**
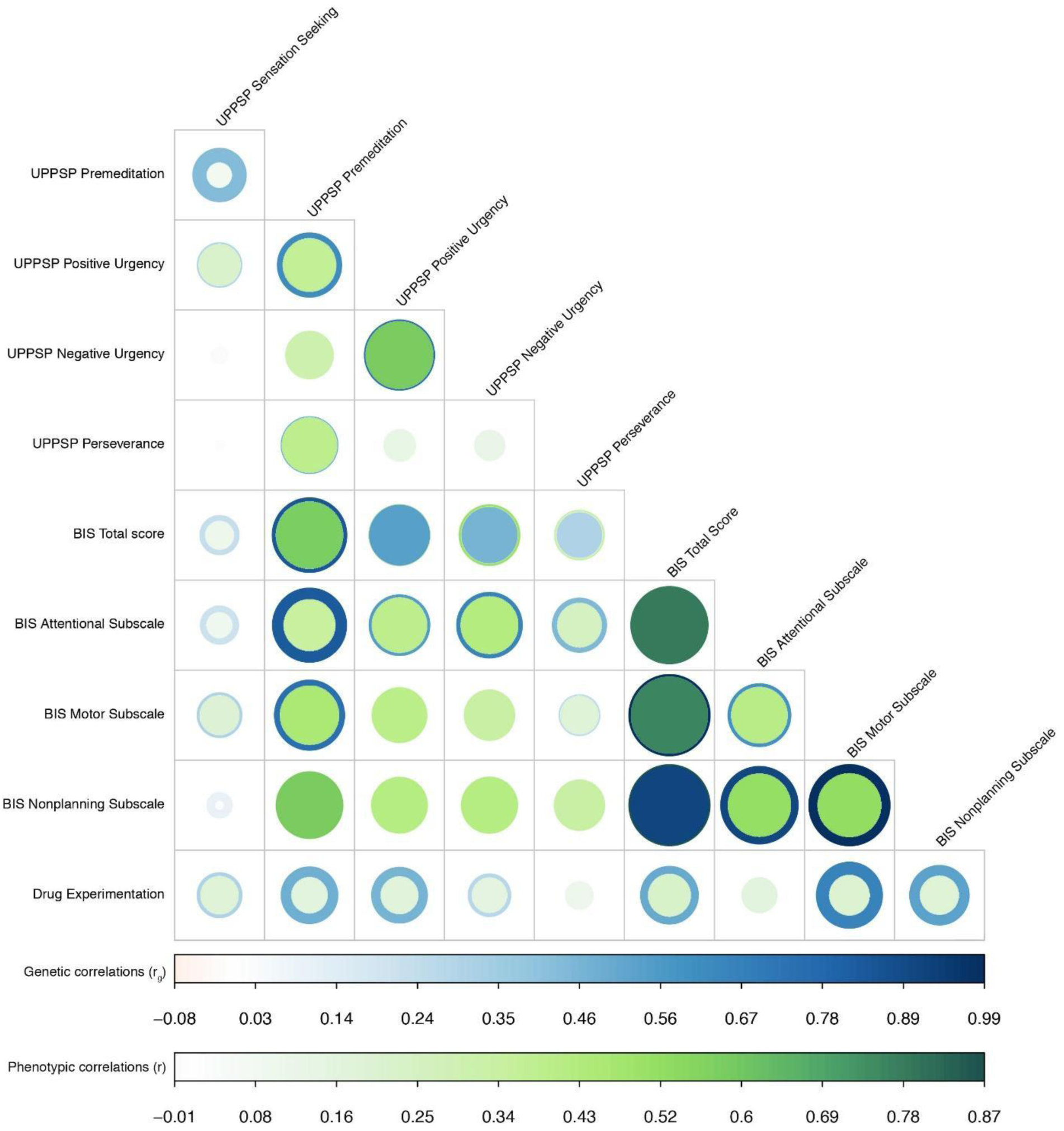
Phenotypic (r, in green) and genetic (r_g_, in blue) correlations between UPPSP and BIS subscales; corresponding p-values are shown in **Supplementary Tables 20 & 21**. The width and color gradient of the circles indicate the strength of the correlation.

With regards to other putatively distinct measures of impulsivity, only impulsivity as quantified by BIS (Total Score and the Nonplanning subscale) showed a positive genetic correlation with delay discounting (i.e., a measure of capacity to delay gratification; r_g_ = 0.49 & 0.72, p < 1.59 x 10^-2^) as measured by the Monetary Choice Questionnaire (48), further emphasizing that impulsivity has multiple distinct facets.

IPT has been established as an important contributor to drug use vulnerability (8). Using multiple independent cohorts, we found modest associations between IPTs and substance use traits (see **Supplementary Figure 25**). Similarly IPT, as measured by BIS (Total Score, Motor, Nonplanning) and UPPSP (Positive Urgency only), showed positive genetic correlations with our measure of drug experimentation (r_g_ = 0.50-0.65, p < 1.09 x 10^-2^; **Supplementary Table 19**). UPPSP subscales (Premeditation, Positive Urgency) and all BIS subscales (except attention) showed positive genetic correlations with lifetime cannabis and tobacco use (r_g_ = 0.46-0.69, p = 1.22 x 10^-3^), suggesting that IPT and drug use share a common genetic basis.

With regards to alcohol use phenotypes, we observed a positive genetic correlation between UPPSP subscales (Sensation Seeking, Premeditation and Positive Urgency) and alcohol consumption (r_g_ = 0.30-0.44, p = 8.50 x 10^-3^), and between UPPSP Premeditation and alcohol dependence (r_g_ = 0.69, p < 1.25 x 10^-2^). Intriguingly, neither Sensation Seeking nor Positive Urgency were genetically correlated with DSM-IV diagnosis of alcohol dependence (49), suggesting a distinction between frequency of use and problem use. Conversely, measures from BIS (Total Score, Nonplanning) showed a positive genetic correlation with alcohol dependence (r_g_ = 0.67 & 0.70, p < 2.00 x 10^-3^) but not alcohol consumption (r_g_ = 0.25, p < 3.65 x 10^-2^). Furthermore, we observed positive genetic correlations between all AUDIT scores (AUDIT Total Score, and the domains pertaining to alcohol consumption and problematic alcohol use: AUDIT-C, AUDIT-P) and several measures from the UPPSP questionnaire, including Sensation Seeking (r_g_ = 0.40-0.52, p < 6.80 x 10^-5^), Premeditation (r_g_ = 0.44-0.59, p < 5.33 x 10^-3^), and Positive Urgency (r_g_ = 0.26-0.54, p < 1.32 x 10^-2^). Surprisingly, only AUDIT-P (but not AUDIT-C or AUDIT Total Score) showed a significant positive genetic correlation with UPPSP Negative Urgency (r_g_ = 0.34, p = 3.22 x 10^-3^). On the contrary, AUDIT-total and AUDIT-C (but not AUDIT-P) showed a positive correlation with the UPPSP Perseverance subscale (r_g_ = 0.29, p < 3.14 x 10^-3^). Similarly, all AUDIT dimensions showed positive genetic correlations with the BIS Nonplanning subscale (rg = 0.28 & 0.49, p < 1.74 x 10^-2^), whereas only BIS Total Score and BIS Attentional subscales showed positive genetic correlations with AUDIT-P (r_g_ = 0.34 & 0.48, p = 1.05 x 10^-2^). Overall, the correlations with alcohol consumption and dependence could suggest that different facets of IPT influence different temporal stages on the path from alcohol consumption to dependence.

With regards to other personality measures, UPPSP Sensation Seeking and BIS Motor subscales showed a positive genetic correlation with extraversion (r_g_ = 0.53-0.56, p < 8.36 x 10^-3^). In addition, UPPSP Sensation Seeking showed a negative genetic correlation with neuroticism (r_g_ = -0.30, p = 1.87 x 10^-3^), whereas UPPSP (positive and Negative Urgency) and BIS (Total Score, Attentional) showed positive genetic correlations with neuroticism (r_g_ = 0.30-0.60, p< 1.02 x 10^-2^).

Considering our measure of drug experimentation, we found positive correlations with other substance use phenotypes, including alcohol consumption (r_g_ = 0.48, p = 6.15 x 10^-7^), AUDIT scores (AUDIT Total Score, AUDIT-C, AUDIT-P; r_g_ = 0.55-0.59, p < 4.03 x 10^-7^), alcohol dependence (r_g_ = 0.68, p = 3.02 x 10^-5^), lifetime tobacco (r_g_ = 0.82, p= 1.61 x 10^-12^) and cannabis smoking (r_g_ = 1.02, p = 7.65 x 10^-12^).

Impulsivity is a core symptom of ADHD (3, 22). We found that UPPSP Premeditation and Positive Urgency and all BIS subscales showed positive genetic correlations with ADHD (rg = 0.43 & 0.30, p < 8.69 x 10^-3^; r_g_ = 0.44-0.51, p < 9.37 x 10^-3^). We also identified a positive genetic correlation between drug experimentation and ADHD (_rg_ = 0.28, p = 1.23 x 10^-3^).

Impulsivity may also play a role in other psychiatric disorders (3). For example, we found that, BIS Total Score and UPPSP positive and Negative Urgency showed a positive genetic correlation with depressive symptoms (r_g_ = 0.33 & 0.30 & 0.45, p < 7.90 x 10^-3^). Similarly, BIS Total Score, BIS attention and UPPSP Negative Urgency showed negative genetic correlations with subjective wellbeing (r_g_ = -0.33 & -0.49 & - 0.48, p < 1.72 x 10^-2^).

Lastly, BIS (Total Score, Nonplanning) and UPPSP Negative Urgency showed negative genetic correlations with educational variables (r_g_ = -0.30-0.50 & -0.39 & -0.29, p < 1.37 x 10^-2^), whereas UPPSP Sensation Seeking and UPPSP Perseverance showed positive genetic correlations with educational measures (r_g_ = 0.16-0.39, p < 6.43 x 10^-3^). Drug experimentation showed a negative genetic correlation with years of education (r_g_ = -0.14, p = 1.39 x 10^-2^).

## DISCUSSION

We have performed the largest GWAS of self-reported impulsive personality traits to date. We identified an association between SNPs in *CADM2* and Sensation Seeking and drug experimentation. In addition, we identified an association between variants in *CACNA1I* and Negative Urgency. This correlation was further supported by single variant, gene- and transcriptome-based analyses. Impulsive personality traits and drug experimentation were modestly heritable (5-11%). This study is the first to provide evidence that the construct of impulsivity can be genetically separated into distinct components. We showed that impulsive personality traits are highly associated with both substance use involvement and ADHD at the genetic level, suggesting impulsivity is an endophenotype contributing to these psychiatric conditions.

High impulsivity depends on a neural network that includes the ventral striatum (subsuming the nucleus accumbens) with top-down control from prefrontal cortical regions, and is modulated by monoamine neurotransmitters including dopamine and serotonin (50). In contrast to various candidate gene studies, in the present analysis polymorphisms in genes associated brain monoamine function (19, 51) were not associated without specifying a threshold (i.e. P > 5 x 10^-8^) with self-reported impulsivity traits. Instead, we identified signal in the gene Cell adhesion molecule 2 (*CADM2*, aka *SynCAM2*), which encodes a mediator of synaptic signaling. Independent GWASs have previously identified significant associations between risky or impulsive behaviors [risk-taking (35), risk-tolerance, automobile speeding propensity, number of sexual partners (36); high alcohol consumption (39); high AUDIT scores (38); cannabis lifetime use (40)] and multiple loci in *CADM2*. Many of these variants are expression quantitative trait loci in multiple tissues, including the brain (i.e. basal ganglia, cerebellum, hippocampus; see **Supplementary Table 9**). Our lead SNPs (rs139528938 and rs2163971), which were associated with Sensation Seeking and drug experimentation, respectively, are highly correlated with one another (r^2^ = 0.68, r^2^ = 0.57) and with a previously identified lead *CADM2* SNP (rs57401290, r^2^ = 0.57) that showed genome-wide significant associations in the UK Biobank study with self-reported risk-taking propensity (rs57401290: p = 5.3 × 10^-9^), and nominal associations with number of sexual partners (rs57401290: p = 6.0 × 10^-7^) and number of children (rs57401290: p = 6.2 × 10^-7^) (36). C*ADM2* is expressed throughout the brain, and modulates synapse assembly (52). *CADM2* encodes an immunoglobulin-domain-containing adhesion protein that spans the synaptic cleft and induces excitatory synapses. CADMs are highly expressed in cholinergic interneurons in the striatum, whose arborization may affect striatal circuits (53). Intriguingly, a closely related synapse-organizing protein, CADM1, has been implicated in drug addiction (53, 54) and appetitive behaviors characterized by elevated impulsivity [i.e. food intake, obesity (55)], for which subcortical circuits are essential.

Impulsivity is increasingly recognized as a phenotypically heterogeneous construct, and our LDSC genetic correlation analyses provide novel genetic evidence to support this view (56, 57). The current data support the idea that the diverse impulsivity traits (measured by BIS, UPPSP, delay discounting questionnaires) may be governed by both overlapping and distinct genetic substrates (**Figure 2**, **Figure 3**). Particularly, Sensation Seeking did not exhibit genetic associations with UPPSP or BIS dimensions. Although Sensation Seeking is considered an impulsivity-related trait within the UPPS measure, these data suggest that the preference for highly stimulating experiences is genetically distinct from all the other BIS and UPPSP subscales, and it is consistent with earlier phenotypic observations (20).

Other personality traits, such as extraversion and neuroticism (58, 59), have sometimes been considered closely related to impulsive personality traits (10). Our genetic correlation analyses provide more evidence for some of these associations; Sensation Seeking scores showed a positive genetic correlation with extraversion and a negative genetic correlation with neuroticism, whereas UPPSP (positive and Negative Urgency) and BIS (total, attention) showed positive genetic correlations with neuroticism (60).

We examined the relevance of UPPSP and BIS facets of impulsivity as surrogates for substance use disorders. Epidemiological studies have also shown that impulsivity is elevated in drug using individuals; however, such studies are based on phenotypic correlations and cannot therefore identify causality mechanisms (i.e. impulsivity could either precede or result from drug use). By studying impulsivity in research participants with low rates of drug use we were able to study normal variation in IPTs without the confounding influence of drug use. We found that impulsivity and substance use have a common genetic etiology (see **Supplementary Figure 25**), demonstrating that forms of impulsivity meet the first criteria necessary to be considered an endophenotype for substance use disorders (61, 62).

We have also shown that different facets of IPTs may mediate use of specific drug types, as has been found in phenotypic correlations between Motor impulsivity and stimulant abuse (24, 63), cognitive impulsivity for cocaine and heroin use (64), and different UPPS measures predicting different aspects of alcohol use [alcohol consumption, misuse and dependence; (65)]. For example, we have shown that some UPPSP (lack of Premeditation) and BIS (Total Score, Nonplanning) measures were genetically correlated with DSM-IV diagnosis of alcohol dependence, whereas some UPPSP measures (lack of Premeditation, Positive Urgency, Sensation Seeking) correlated with alcohol consumption and AUDIT Total Score.

Similarly, other studies have reported that individuals with ADHD exhibit elevated impulsive personality traits (65, 66) and high rates of drug use (67); here we showed that all BIS measures and some UPPSP subscales (lack of Premeditation, Positive Urgency), in addition to drug experimentation, showed positive genetic correlations with ADHD, demonstrating that impulsivity could also serve as an endophenotype for ADHD (68). Finally, we identified positive genetic correlations of BIS total and UPPS positive and Negative Urgency with depressive symptoms. These observations illustrate the role of impulsivity in multiple psychiatric diagnoses, consistent with the NIMH Research Domain Criteria (**RDoC**) approach (69).

Our study is not without limitations. Our research participants showed generally low levels of IPT scores in absolute terms. It is not known whether the genetic correlations observed here would extend to cohorts with higher levels of IPTs. It is also not known whether the correlations would apply to samples reporting higher rates of drug use, although the lack of extensive drug use in our sample minimizes the chance that prolonged exposure to drugs of abuse increased impulsivity levels [i.e. (70)]. Another limitation is that we measured self-reported impulsive personality, and did not include behavioral measures (71–76). Self-report measures can provide evidence of ‘trait’ impulsivity (stable personality characteristic), whereas performance-based tests may measure aspects of ‘state’ impulsivity (influenced by environmental variables). Although self-report measures are useful, this methodology is limited as it may be influenced by subjective bias [e.g. less insight of the inhibitory control deficits in drug abusers (77) or by the subject’s state (78)]. Also, impulsivity traits may predispose individuals to take drugs, which in turn may lead to further (state) impulsivity, manifested in a failure to control drug taking behavior (79). Nonetheless, these research participants show low rates of drug use, minimizing the possibility that individual differences were a result, rather than a cause, of drug use. Moreover, impulsivity changes across the lifespan; although there is empirical support for impulsivity being a stable trait (57), our findings may not generalize to younger populations. As previously reviewed in a meta-analysis, total genetic effects across various impulsivity traits were found to be important for all ages, but appeared to be strongest in children (15, 80).

Our results indicate that impulsive personality traits are influenced by numerous genetic variants, and dissecting the genetics of impulsivity would likely benefit from studies with even larger sample sizes. Variants in the *CADM2* gene, implicated in recent GWAS of risk-associated traits [i.e. (36, 39, 41)], are associated with Sensation Seeking, and nominally with drug experimentation. We are currently establishing mutant mouse and rat lines of this gene to dissect the molecular events that underlie this trait. Altogether, this study is the first to demonstrate a role for common genetic contribution to individual differences in impulsive personality traits, as well as genetic overlap with human psychopathology, particularly ADHD. It also features associations with both the degree of drug experimentation, and with later stages of alcohol use. Our approach shows how genetic studies that measure multiple impulsivity traits can be used to gain insight into the biology of neuropsychiatric diseases.

## ACKNOWLEDGEMENTS AND DISCLOSURES

We would like to thank the 23andMe research participants and employees for making this work possible. J.M.’s contributions were partially supported by the Peter Boris Chair in Addictions Research and he discloses that he is a principal in BEAM Diagnostics, Inc. S.S-R was supported by the Frontiers of Innovation Scholars Program (FISP; #3-P3029), the Interdisciplinary Research Fellowship in NeuroAIDS (IRFN; MH081482) and a pilot award from DA037844. Pierre Fontanillas, Sarah L. Elson, and members of the 23andMe Research Team are employees of 23andMe, Inc. The opinions and assertions expressed herein are those of the author(s) and do not necessarily reflect the official policy or position of the Uniformed Services University or the Department of Defense.

## DATA AVAILABILITY

The full GWAS summary statistics will be made available through 23andMe to qualified researchers under an agreement that protects participant privacy. Interested researchers should visit https://research.23andme.com/collaborate/#publication to learn more and to apply to access the data.

**Table 1.**
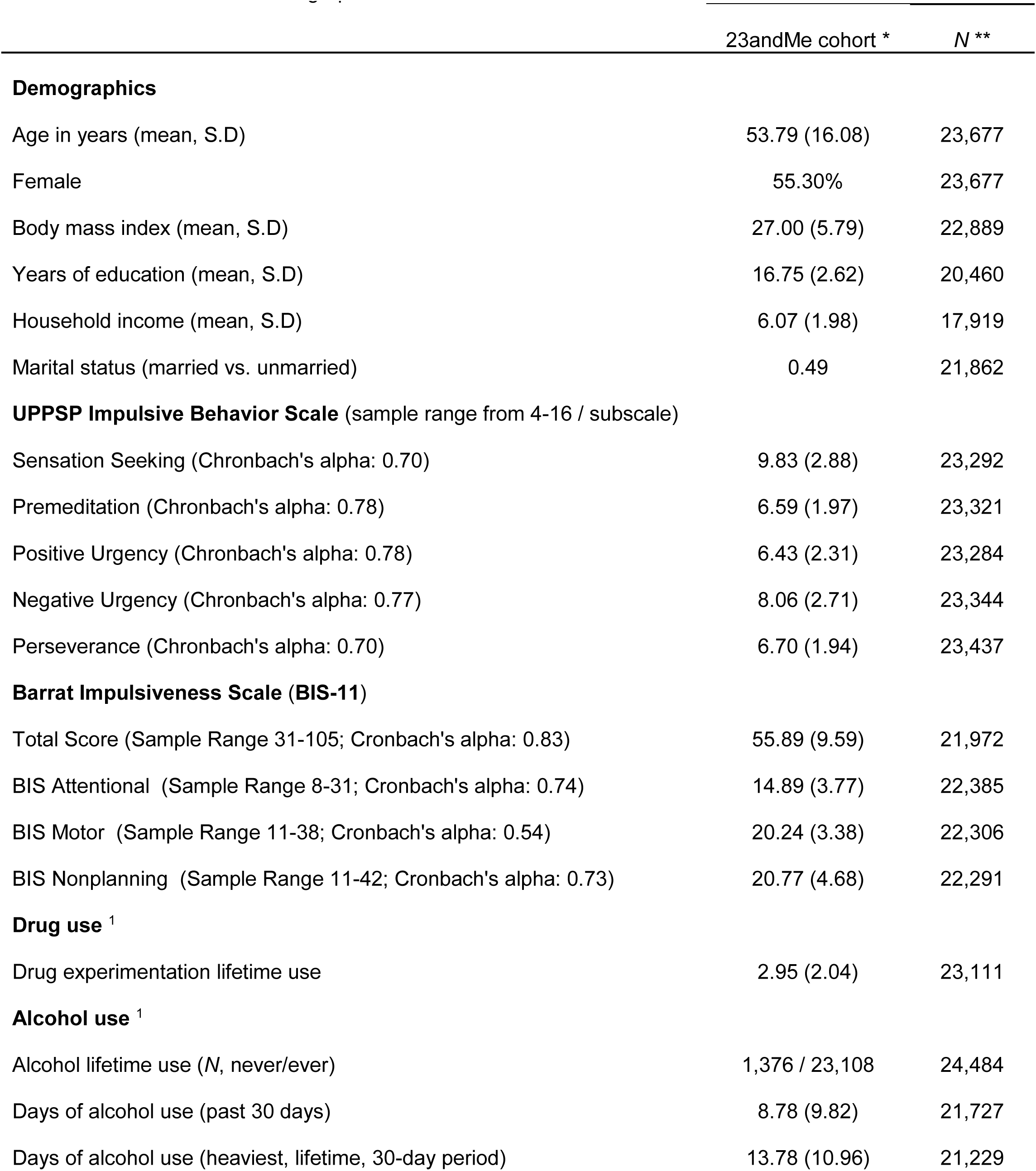
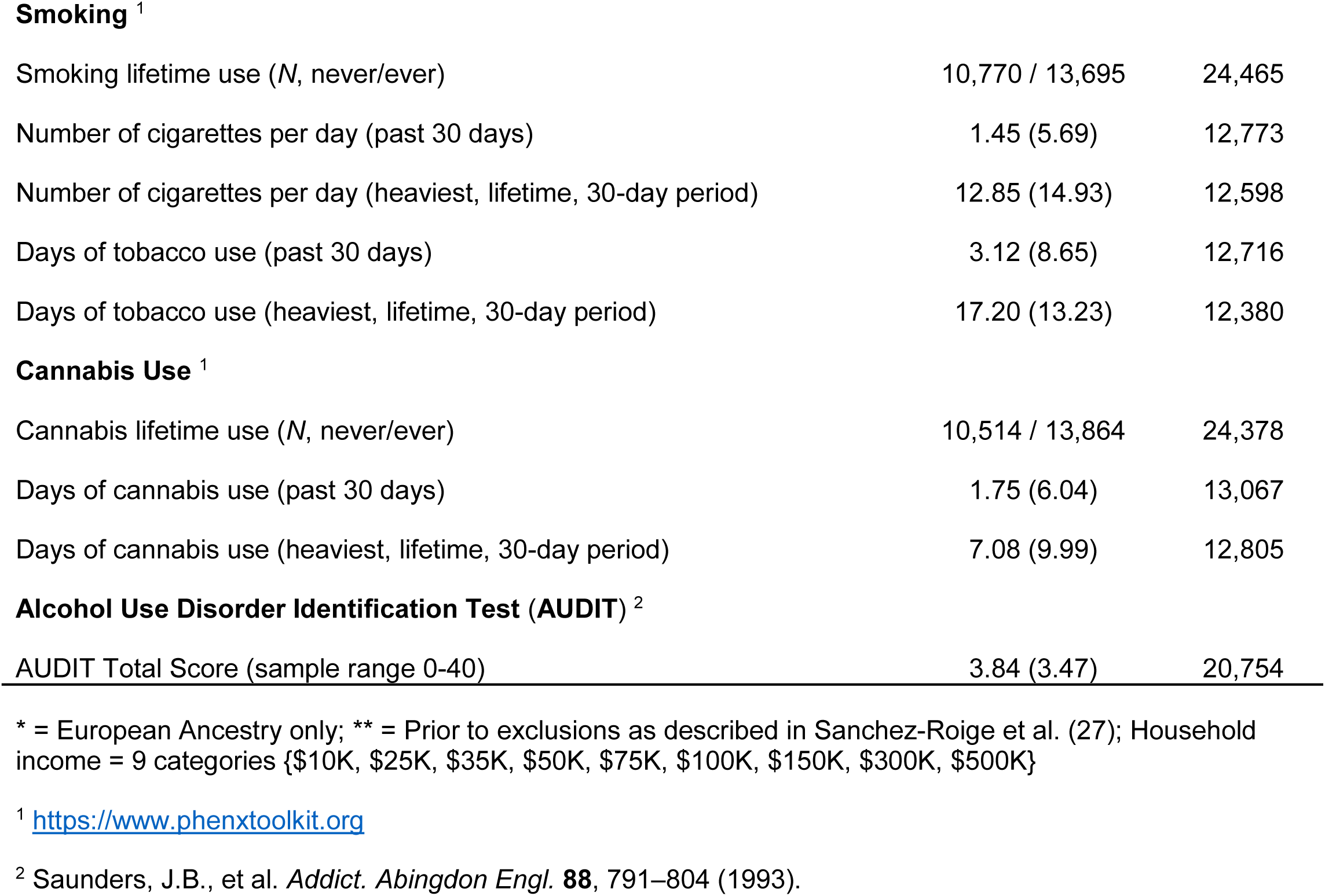
This table shows demographic characteristics of the 23andMe coh<u>ort</u>23andMe cohort

|  | 23andMe cohort * | N ** |
| --- | --- | --- |
| **Demographics** |  |  |
| Age in years (mean, S.D) | 53.79 (16.08) | 23,677 |
| Female | 55.30% | 23,677 |
| Body mass index (mean, S.D) | 27.00 (5.79) | 22,889 |
| Years of education (mean, S.D) | 16.75 (2.62) | 20,460 |
| Household income (mean, S.D) | 6.07 (1.98) | 17,919 |
| Marital status (married vs. unmarried) | 0.49 | 21,862 |
| **UPPSP Impulsive Behavior Scale** (sample range from 4-16 / subscale) |  |  |
| Sensation Seeking (Chronbach's alpha: 0.70) | 9.83 (2.88) | 23,292 |
| Premeditation (Chronbach's alpha: 0.78) | 6.59 (1.97) | 23,321 |
| Positive Urgency (Chronbach's alpha: 0.78) | 6.43 (2.31) | 23,284 |
| Negative Urgency (Chronbach's alpha: 0.77) | 8.06 (2.71) | 23,344 |
| Perseverance (Chronbach's alpha: 0.70) | 6.70 (1.94) | 23,437 |
| **Barrat Impulsiveness Scale (BIS-11)** |  |  |
| Total Score (Sample Range 31-105; Cronbach's alpha: 0.83) | 55.89 (9.59) | 21,972 |
| BIS Attentional (Sample Range 8-31; Cronbach's alpha: 0.74) | 14.89 (3.77) | 22,385 |
| BIS Motor (Sample Range 11-38; Cronbach's alpha: 0.54) | 20.24 (3.38) | 22,306 |
| BIS Nonplanning (Sample Range 11-42; Cronbach's alpha: 0.73) | 20.77 (4.68) | 22,291 |
| **Drug use <sup>1</sup>** |  |  |
| Drug experimentation lifetime use | 2.95 (2.04) | 23,111 |
| **Alcohol use <sup>1</sup>** |  |  |
| Alcohol lifetime use (N, never/ever) | 1,376 / 23,108 | 24,484 |
| Days of alcohol use (past 30 days) | 8.78 (9.82) | 21,727 |
| Days of alcohol use (heaviest, lifetime, 30-day period) | 13.78 (10.96) | 21,229 |
| **Smoking** <sup>1</sup> |  |  |
| Smoking lifetime use (*N*, never/ever) | 10,770 / 13,695 | 24,465 |
| Number of cigarettes per day (past 30 days) | 1.45 (5.69) | 12,773 |
| Number of cigarettes per day (heaviest, lifetime, 30-day period) | 12.85 (14.93) | 12,598 |
| Days of tobacco use (past 30 days) | 3.12 (8.65) | 12,716 |
| Days of tobacco use (heaviest, lifetime, 30-day period) | 17.20 (13.23) | 12,380 |
| **Cannabis Use** <sup>1</sup> |  |  |
| Cannabis lifetime use (*N*, never/ever) | 10,514 / 13,864 | 24,378 |
| Days of cannabis use (past 30 days) | 1.75 (6.04) | 13,067 |
| Days of cannabis use (heaviest, lifetime, 30-day period) | 7.08 (9.99) | 12,805 |
| **Alcohol Use Disorder Identification Test (AUDIT)** <sup>2</sup> |  |  |
| AUDIT Total Score (sample range 0-40) | 3.84 (3.47) | 20,754 |

\* = European Ancestry only; \*\* = Prior to exclusions as described in Sanchez-Roige et al. (27); Household income = 9 categories {\$10K, \$25K, \$35K, \$50K, \$75K, \$100K, \$150K, \$300K, \$500K}
<sup>1</sup> https://www.phenxtoolkit.org
<sup>2</sup> Saunders, J.B., et al. *Addict. Abingdon Engl.* **88**, 791–804 (1993).

